# Explainable AI for end-to-end pathogen target discovery and molecular design

**DOI:** 10.64898/2026.02.27.708593

**Authors:** Lucía Jiménez-Castro, Dolores Fernández-Ortuño, Alejandro Pérez-García, Álvaro Polonio

## Abstract

Drug discovery is limited by target identification, a major bottleneck in antimicrobial development and in combating emerging fungicide resistance. We present APEX (Attention-based Protein EXplainer), an explainable AI framework for cross-species, proteome-scale target discovery and pocket-guided molecular design. APEX combines ESM-2 embeddings, GAT, and a MLP to train pathogen-specific essentiality predictors (APEX-Tar) alongside a universal druggability model (APEX-Drug). Attention maps and GNNExplainer-derived subgraphs highlight residues driving predictions, enabling direct conditioning of structure-based diffusion models for inhibitor generation. APEX-Tar identifies key residues in known fungal targets and nominates new candidates, including adenylosuccinate lyase (ADSL) in fungi and the bacterial adhesin YadV. APEX-Drug recapitulates established fungicide binding sites, guides the design of ADSL inhibitors exploiting a fungal-specific active-site residue, and uncovers in YadV a previously undescribed pocket distinct from known pilicide sites. Together, APEX provides a kingdom-agnostic, explainable pipeline for target prioritization and guided molecular design, accelerating the search for next-generation antimicrobials.

---

The search for new therapeutic agents is increasingly constrained by one persistent bottleneck: identifying the right targets. Despite major advances in chemical synthesis and high-throughput screening, target discovery remains slow, fragmented, and heavily dependent on resource-intensive experimental workflows^1,2^. In 2024, only a small number of approved drugs engaged new mechanisms of action, underscoring how difficult it remains to expand the druggable genome^2^. The absence of integrated computational pipelines capable of systematically identifying targetable proteins and designing inhibitors further compounds this limitation^3,4^.

This bottleneck is especially damaging in antimicrobial resistance and crop protection. Antimicrobial resistance is projected to cause tens of millions of deaths in the coming decades^5^, yet antibiotic development suffers from both a scarcity of candidates and minimal mechanistic innovation^6^. Between 2014 and 2021, only 18 new antibiotics were approved, just two with truly novel modes of action^7^. Without new strategies to uncover unexplored targets, multidrug-resistant pathogens threaten to render common infections untreatable^8^. Agriculture faces parallel pressures: plant-pathogenic fungi drive severe global crop losses^9,10^, and reliance on a narrow set of fungicide targets has accelerated the emergence of cross-resistant strains^11,12^.

These challenges arise at a moment when artificial intelligence is transforming biological research. Massive investments in AI-driven biology have enabled breakthroughs in protein structure prediction (e.g., AlphaFold)^13,14^ and protein language modeling (e.g., ESM)^15^, opening access to biological and chemical spaces previously unreachable through traditional experimentation. Transformer architectures naturally extend to protein sequences, building on advances in natural language processing^16,17^. Large language models such as GPT and BERT learn contextual representations through self-supervised training, and the same principles (self-attention, masked-token prediction, and large-scale pretraining) now underpin protein language models (PLMs). Models like ESM-2 capture structural motifs, functional domains, and evolutionary patterns, enabling accurate structure prediction and functional annotation^18–20^.

As model complexity increases, interpretability becomes essential. Black-box predictions offer limited mechanistic insight, and post-hoc explainability tools such as SHAP^21^ and LIME^22^ struggle with scalability, correlated features, and causal grounding^23^. In contrast, Graph Attention Networks (GATs) ^24^ provide intrinsic interpretability through learned attention weights that highlight influential residues and interactions ^25,26^. Complementary methods like GNNExplainer^27^ identify minimal subgraphs that drive predictions, producing explanations aligned with protein structure and biochemistry. Meanwhile, drug discovery increasingly relies on generative modeling. Classical de novo design, rule-based assembly or virtual screening, explore only a narrow chemical space. Diffusion models^28,29^ now offer a powerful alternative: by learning probability distributions over 3D molecular geometries, they generate realistic, symmetry-aware structures through iterative denoising^30^. When conditioned on protein pockets, these models can produce ligands tailored to specific binding sites, shifting hit discovery from empirical screening to systematic computational design^29,31^.

Here we introduce APEX (Attention-based Protein EXplainer), an AI pipeline for antimicrobial target prioritization and inhibitor design. APEX integrates ESM-2 embeddings, predicted contact maps and GATs in two models: APEX-Tar, which identifies essential or virulence-associated pathogen proteins, and APEX-Drug, a universal druggability classifier. At proteome scale, APEX jointly ranks targets by essentiality and druggability. Its interpretability layer maps functional and druggable regions using attention weights and GNNExplainer subgraphs, guiding structure-based diffusion models towards pocket-specific ligand generation. Together, APEX links target discovery to pocket-conditioned molecular design

## Results

### An end-to-end pipeline for rational design of pathogen inhibitors

Our workflow is built around APEX, a graph neural network framework that integrates ESM-2 protein embeddings with explainable AI (XAI) to identify which pathogen proteins matter and where they can be most effectively targeted. By linking these insights to structure-based diffusion models, the pipeline forms a continuous path from target prioritization to the generation of site-tailored small molecules (Figure 1). The process begins with curated training datasets (Figure 1a). APEX-Tar learns to distinguish essential pathogen proteins from non-essential ones, while APEX-Drug is trained on a human dataset separating druggable from non-druggable proteins. Both models share a unified architecture combining ESM-2 embeddings, GAT and multilayer perceptron classifiers (Figure 1b). This design allows APEX-Tar to adapt to different pathogen groups while APEX-Drug remains a general predictor of druggability. Once trained, the models are deployed across the pathogen proteome (Figure 1c). Each protein receives a composite score reflecting its functional relevance and its likelihood of being druggable. High-scoring proteins then undergo an interpretability analysis (Figure 1d): APEX-Tar highlights key residues for essentiality, whereas APEX-Drug identifies structurally coherent regions with high druggability potential. These XAI-defined pockets serve as direct inputs to diffusion-based molecular generation (Figure 1e). The structural constraints guide the model to produce small molecules tailored to the predicted binding sites, completing an automated pipeline that moves seamlessly from target selection to the proposal of rational pathogen inhibitors.

**Figure 1.**
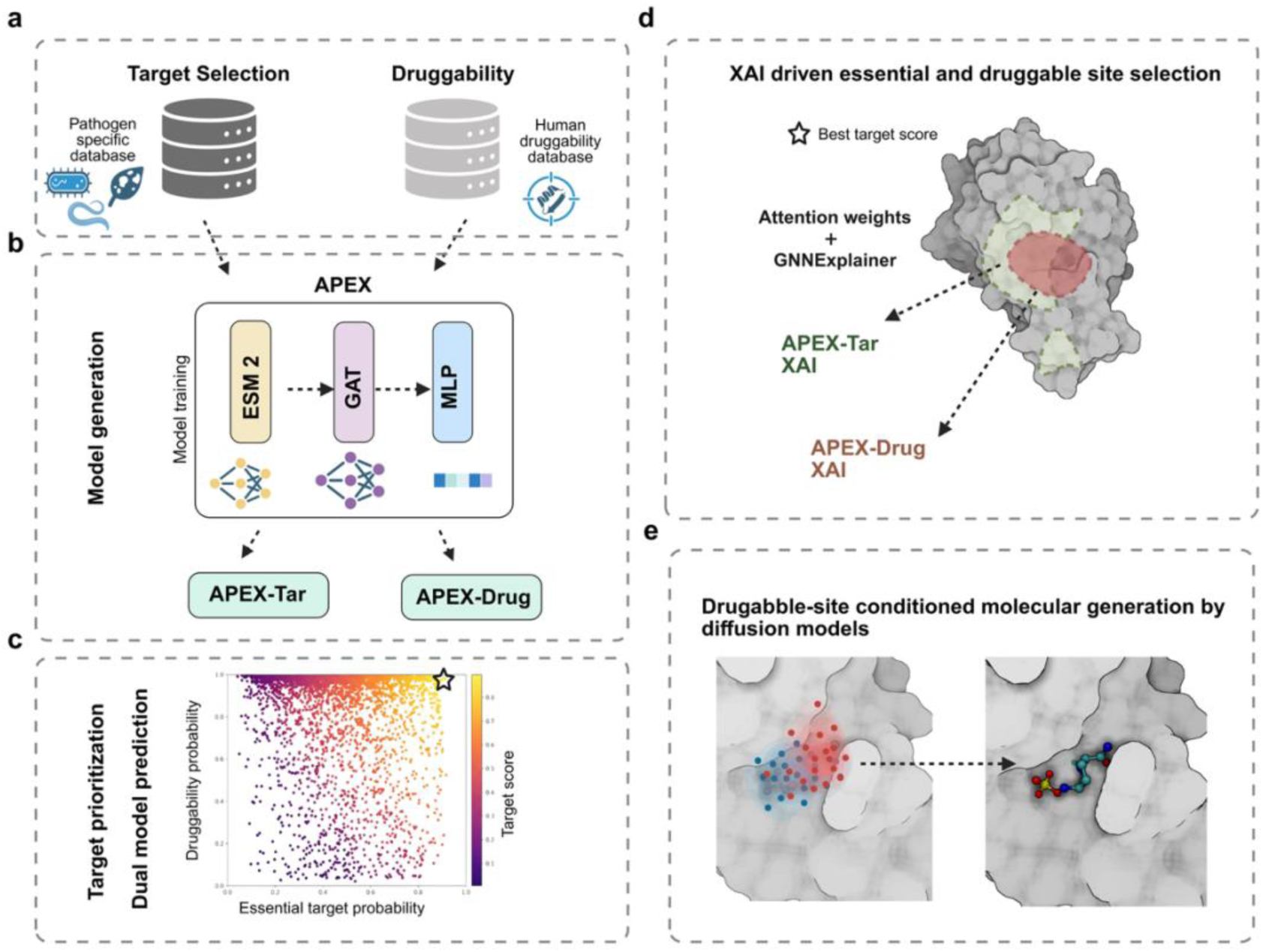
End-to-end pipeline for AI-driven pathogen target discovery and inhibitor design. (a) Training datasets used for model development: pathogen-specific essential and virulence sets for APEX-Tar, and a human druggability dataset for APEX-Drug. (b) Unified architecture integrating ESM-2 embeddings, GAT layers, and an MLP classifier, implemented as task-specific variants for target prediction and druggability assessment. (c) Proteome scale target prioritization by integrating APEX-Tar and APEX-Drug outputs to identify high-confidence protein candidates. (d) XAI-guided site selection using attention weights and GNNExplainer to highlight essential structural regions and druggable pockets. (e) Diffusion-based molecular generation conditioned on predicted druggable sites to produce candidate inhibitors.

### APEX architecture: unifying sequence embeddings and structural features for interpretable protein classification

Identifying therapeutic targets requires solving two complementary challenges: distinguishing disease-relevant proteins and determining whether those proteins contain druggable sites. To address both tasks, we developed APEX, a unified architecture trained on task-specific datasets but sharing a common representational backbone. APEX consists of three components. First, graphs are constructed from ESM-2 embeddings and contact probability maps (Figure 2a). Residues predicted to be proximal are connected, producing a topology that integrates evolutionary and structural information. Second, a two-layer GAT performs attention-based message passing, followed by global max pooling and a MLP classifier (Figure 2b). This architecture transforms 1,280-dimensional residue embeddings into graph-level representations via a 1,280 → 16 → 1 fully connected network. Third, APEX incorporates built-in interpretability through attention coefficients and GNNExplainer (Figure 2c), enabling residue and subgraph-level explanations.

**Figure 2.**
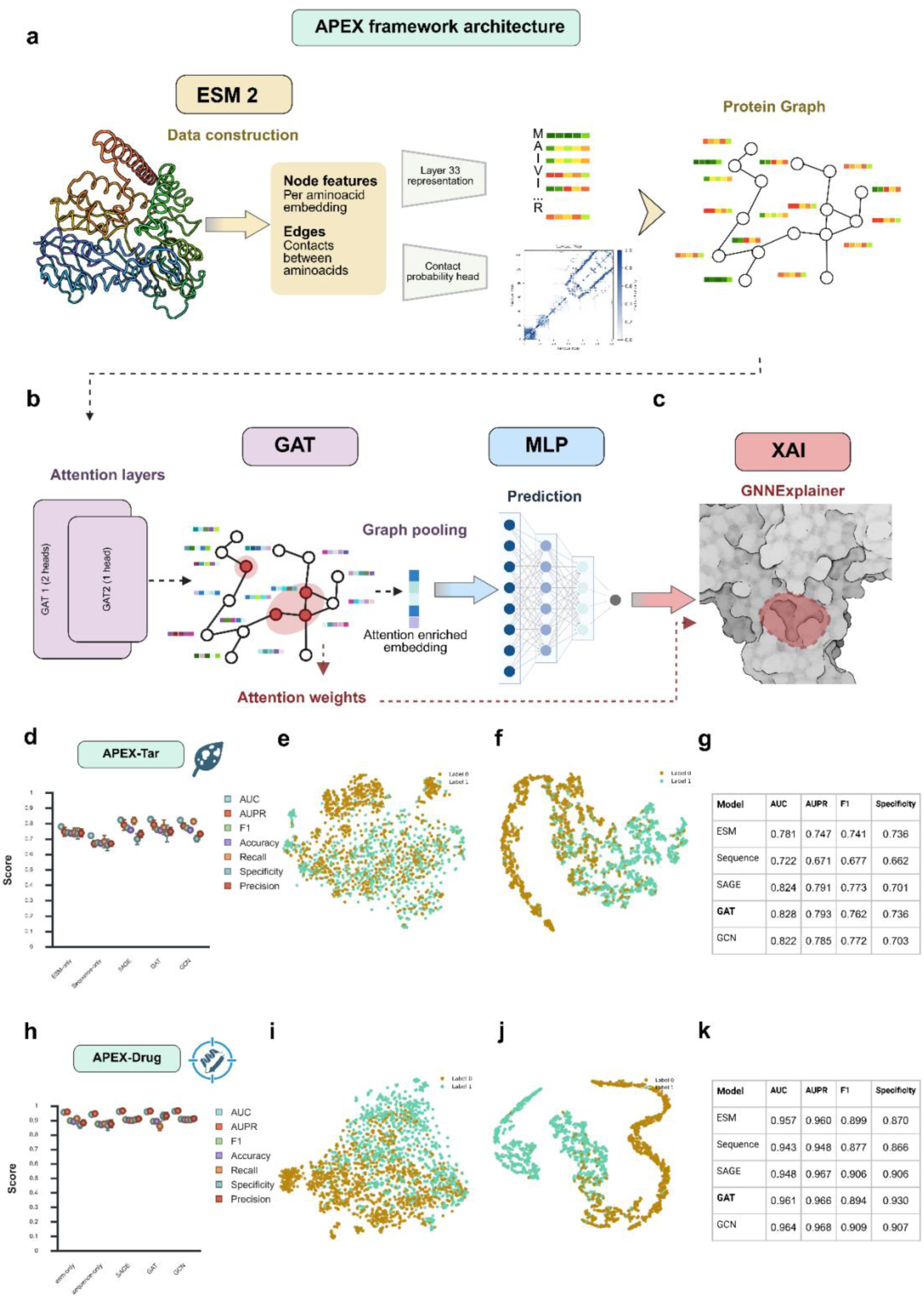
APEX architecture and performance. (a) Graph construction. ESM-2 generates 1,280-dimensional residue embeddings and contact probability maps; contacts above a defined threshold form edges between spatially proximal residues. (b) GAT classifier. Two graph-attention layers perform message passing over the protein graph, followed by global max pooling to obtain a graph-level embedding. A two-layer MLP (1,280 → 16 → 1) produces the final prediction. (c) Explainability. Interpretability is provided through attention-weight visualization and GNNExplainer, which identifies minimal subgraphs most influential for model decisions. (d) Performance of APEX-Tar on fungal pathogenicity prediction across seven metrics using 5-fold cross-validation. (e-f) t-SNE embeddings before and after APEX-Tar training, showing improved separation of pathogenic and non-pathogenic proteins. (g) Model comparison for APEX-Tar. All GNNs (GAT, GCN, GraphSAGE) outperform ESM-only and sequence-only baselines; GAT achieves the highest specificity (0.736). Training set: 1,969 PHI-base fungal proteins. (h) Performance of APEX-Drug on human protein druggability prediction across seven metrics using 5-fold cross-validation. (i–j) t-SNE embeddings before and after APEX-Drug training, showing enhanced separation of druggable versus non-druggable proteins. (k) Model comparison for APEX-Drug. All GNNs achieve high performance (>0.96 AUC), with GAT showing the highest specificity (0.930). Training set: 2,345 ProTar-II human proteins.

Interpretability operates at two scales. Attention coefficients from both GAT layers (with Layer 1 averaged across heads) are aggregated over each residue’s structural neighbors to quantify its importance as an information-integration hub. These length-normalized scores are mapped to sequence positions to generate residue-level profiles. In parallel, GNNExplainer identifies minimal subgraphs supporting individual predictions by optimizing a sparse soft node mask. Explanations are evaluated using insertion–deletion curves, measuring how model confidence changes as top-ranked residues are added or removed.

APEX-Tar was trained on 1,969 fungal proteins (987 pathogenicity factors, 982 non-pathogenic) and evaluated using 5-fold stratified cross-validation (Figure 2d–g). All GNN architectures performed similarly (AUC = 0.822–0.828) and significantly outperformed both the ESM-only (AUC = 0.781) and sequence-only (AUC = 0.722) baselines (p < 0.05). GAT and GCN also showed significant improvements over ESM-only (GAT: Δ = +0.047, p = 0.013; GCN: Δ = +0.041, p = 0.042), demonstrating that explicit residue-contact modeling captures functional information beyond sequence embeddings alone. GAT achieved the highest specificity (0.736), exceeding GraphSAGE (0.701) and GCN (0.703). t-SNE projections revealed limited class separation in raw ESM-2 embeddings (Figure 2e), which improved markedly after GAT training (Figure 2f). To assess potential inflation from residual homology, we identified 1,515 cross-fold homologous pairs (≥50% identity; 0.086% of all pairs). Excluding these proteins yielded AUC = 0.821 versus 0.828 (ΔAUC = −0.006), confirming that performance was not driven by sequence similarity leakage.

APEX-Drug was trained on 2,345 human proteins (1,221 druggable, 1,124 non-druggable). All GNNs achieved high performance (AUC = 0.948–0.964), with GCN slightly outperforming GAT (0.964 vs. 0.961) (Figure 2h–k). All models significantly exceeded the sequence-only baseline (p < 0.05). Improvements over the ESM-only baseline (AUC = 0.957) were consistent but not statistically significant, indicating that druggability is a feature already well captured by ESM-2. GAT achieved the highest specificity (0.930), surpassing GCN (0.907) and GraphSAGE (0.906). Raw ESM-2 embeddings showed substantial class structure (Figure 2i), further refined after GNN training (Figure 2j). Full per-fold metrics are provided in Supplementary File S1. To evaluate cross-organism generalization, the GAT model was applied to 228 experimentally validated druggable proteins from fungal and bacterial pathogens. It correctly classified 219 (96.1%) with a mean predicted druggability of 0.918 ± 0.143, demonstrating that the learned features extend beyond the human proteome (Supplementary File S2).

### Explainable AI in APEX identifies functionally relevant residues in validated

#### Botrytis cinerea targets

To assess whether APEX captures biologically meaningful features, we analyzed attention and GNNExplainer scores for 38 validated proteins of the necrotrophic fungus *Botrytis cinerea*, one of the most important plant pathogens worldwide, spanning both pathogenicity factors (APEX-Tar) and established fungicide targets (APEX-Drug) (Figure 3; Supplementary File S3). Residue-level attention and node-importance from GNNExplainer were compared against InterPro annotations, P2Rank-predicted pockets, and characterized binding sites. All proteins were correctly classified by their respective models. Representative examples are summarized below.

**Figure 3.**
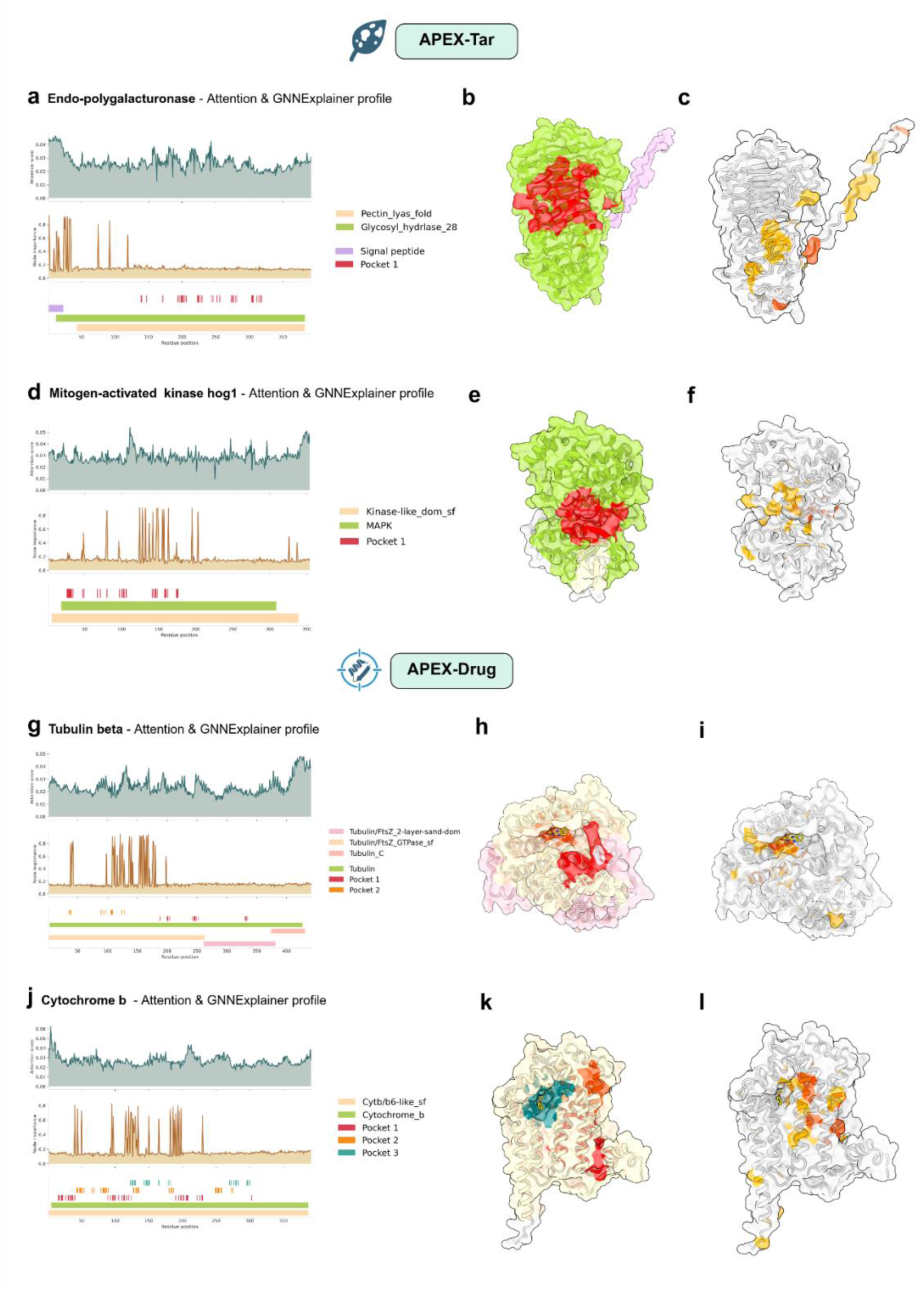
Interpretability of APEX models highlights functionally relevant residues in validated pathogenicity factors and fungicide targets. Rows correspond to analyses of four *B. cinerea* proteins. APEX-Tar panels (a-f): (a–c) endo-polygalacturonase PG1; (d–f) HOG1 MAP kinase. APEX-Drug panels (g-l): (g–i) β-tubulin; (j–l) cytochrome b. The first column (a, d, g, j) shows per-residue attention scores (top, teal) and GNNExplainer importance scores (bottom, orange), with annotated domains and predicted pockets displayed below. The second column (b, e, h, k) present the 3D structural models with functional domains colored and predicted pockets highlighted. The third column (c, f, i, l) maps GNNExplainer-identified important residues onto the structures; the top 10 residues are shown in red-orange, with additional highranking residues in orange. For β-tubulin and cytochrome b, known ligands (benzimidazole and strobilurin, respectively) are shown docked using DiffDock.

##### APEX-Tar: Pathogenicity factors

For endo-polygalacturonase 1, attention was enriched at the N-terminal signal peptide and across regions overlapping Pocket 1 (Figure 3a–b). GNNExplainer highlighted in the signal peptide and pectin lyase fold (Figure 3c), capturing secretion and catalytic features essential for host cell-wall degradation. For Hog1, attention concentrated across the conserved MAPK catalytic domain. GNNExplainer pinpointed residues within the ATP-binding pocket and activation loop (Figure 3d–f), reflecting accurate recognition of the canonical kinase architecture.

##### APEX-Drug: Fungicide targets

For β-tubulin, the target of benzimidazole fungicides, attention spanned all major functional domains (Figure 3g). GNNExplainer focused on residues within the GTPase domain corresponding to colchicine and benzimidazole binding pockets (Figure 3h–i). Docking of carbendazim aligned precisely with these high-importance residues. For cytochrome b, the target of QoI fungicides, attention was distributed across the cytochrome b domain, while GNNExplainer mapped residues to the Qo and Qi sites (Figure 3j–k). The docked azoxystrobin ligand was surrounded by residues identified as important by both interpretability methods (Figure 3l), indicating accurate recognition of the quinone-binding pocket.

Insertion–deletion analysis confirmed GNNExplainer fidelity: adding high-importance residues increased logits, while removing them reduced confidence, confirming that identified residues are causally relevant to predictions. Full insertion-deletion curves and perturbation metrics are provided in Supplementary File S3.

### Target prioritization in *Botrytis cinerea* identifies high-confidence druggable proteins enriched in necrotrophic virulence functions

To identify new antifungal targets, we applied APEX-Tar and APEX-Drug to the complete *B. cinerea* proteome (13,749 proteins). After removing proteins with BLASTp similarity to training datasets or to *Solanum lycopersicum*, 8,046 proteins remained for dual-model prediction. APEX-Tar classified 3,754 proteins (46.7%) as pathogenicity-related, and APEX-Drug identified 2,628 proteins (32.7%) as druggable (P > 0.5), with 1,052 proteins (13%) positive in both models (Supplementary File S4). Applying a stricter combined threshold (P > 0.6) yielded 640 high-priority candidates (Figure 4a).

**Figure 4.**
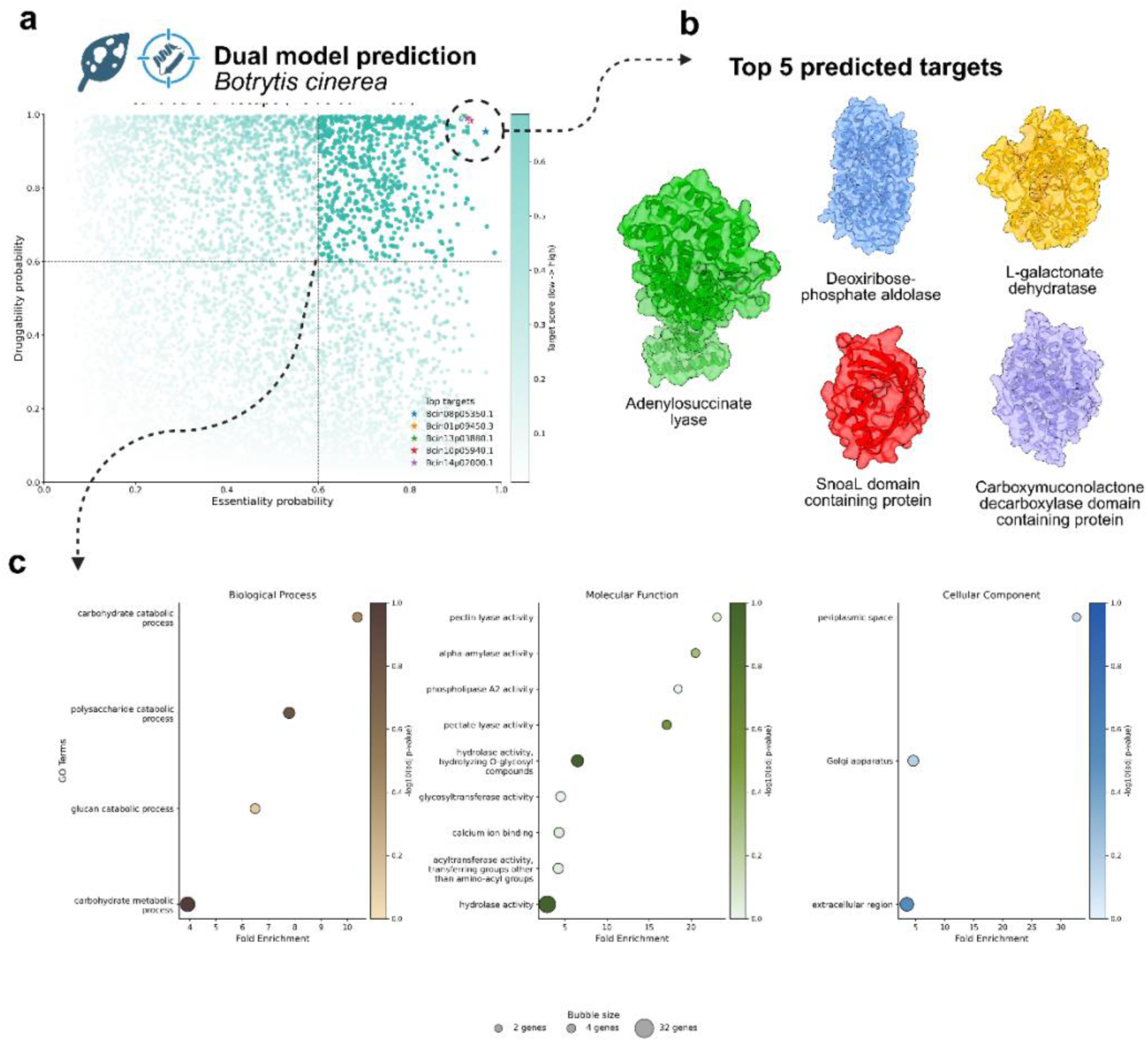
Target prioritization pipeline identifies high-confidence antifungal candidates in *Botrytis cinerea*. (a) Dual-model prediction workflow and outputs. Across 8,046 filtered proteins, APEX-Tar classified 46.7% as pathogenicity-related and APEX-Drug classified 32.7% as druggable, with 13% proteins positive in both models. Applying combined probability thresholds (P > 0.6) yielded 640 high-priority targets. (b) Top five predicted targets (target scores 0.915–0.921): a deoxyribose-phosphate aldolase, a mandelate racemase/muconate lactonizing protein, an adenylosuccinate lyase, a SnoaL-like domain-containing protein, and a carboxymuconolactone decarboxylase-like protein. (c) Functional enrichment of the 640 prioritized targets revealed overrepresentation of biological processes related to carbohydrate and polysaccharide catabolism, molecular functions such as pectin/pectate lyase and glycoside hydrolase activities, and cellular components consistent with active secretion of cell wall-degrading enzymes (extracellular region, Golgi apparatus).

The top five predicted targets (scores 0.921–0.915) corresponded to: a deoxyribose-phosphate aldolase, a mandelate racemase/muconate-lactonizing protein (Bclgd1), an adenylosuccinate lyase (ADSL), a SnoaL-like domain-containing protein, and a carboxymuconolactone decarboxylase-like protein (Figure 4b). All exhibited well-defined catalytic pockets or substrate-binding sites characteristic of druggable protein folds.

Functional enrichment of the 640 high-confidence targets revealed catalytic activity and carbohydrate metabolism as the top overrepresented categories, followed by cell wall and membrane degrading enzymes, glycosyltransferases, and oxidative activities (Figure 4c). Golgi apparatus and extracellular region components were also enriched, together with calcium ion binding and phospholipase A2 activity.

### Explainable AI-guided diffusion modeling enables site-specific inhibitor design

To demonstrate the full end-to-end workflow (from target identification to rational lead generation) we applied our pipeline to adenylosuccinate lyase (ADSL), a central enzyme in de novo purine biosynthesis. Explainability analyses converged on a druggable region located at the oligomeric interface. APEX-Drug attention weights and GNNExplainer importance scores highlighted residues distributed across several predicted pockets, all positioned along the subunit–subunit interfaces required for homotetramer assembly. The highest-scoring residues clustered around the catalytic pocket (Figure 5a). Structural context reinforced these findings: predicted pockets on one dimer aligned with the AMP-docked active site (Figure 5b), confirming geometric and chemical complementarity. Mapping residue importance onto the same dimer (Figure 5c) revealed a compact, high-importance cluster that guided the selection of this pocket for downstream structure-based diffusion modeling.

**Figure 5.**
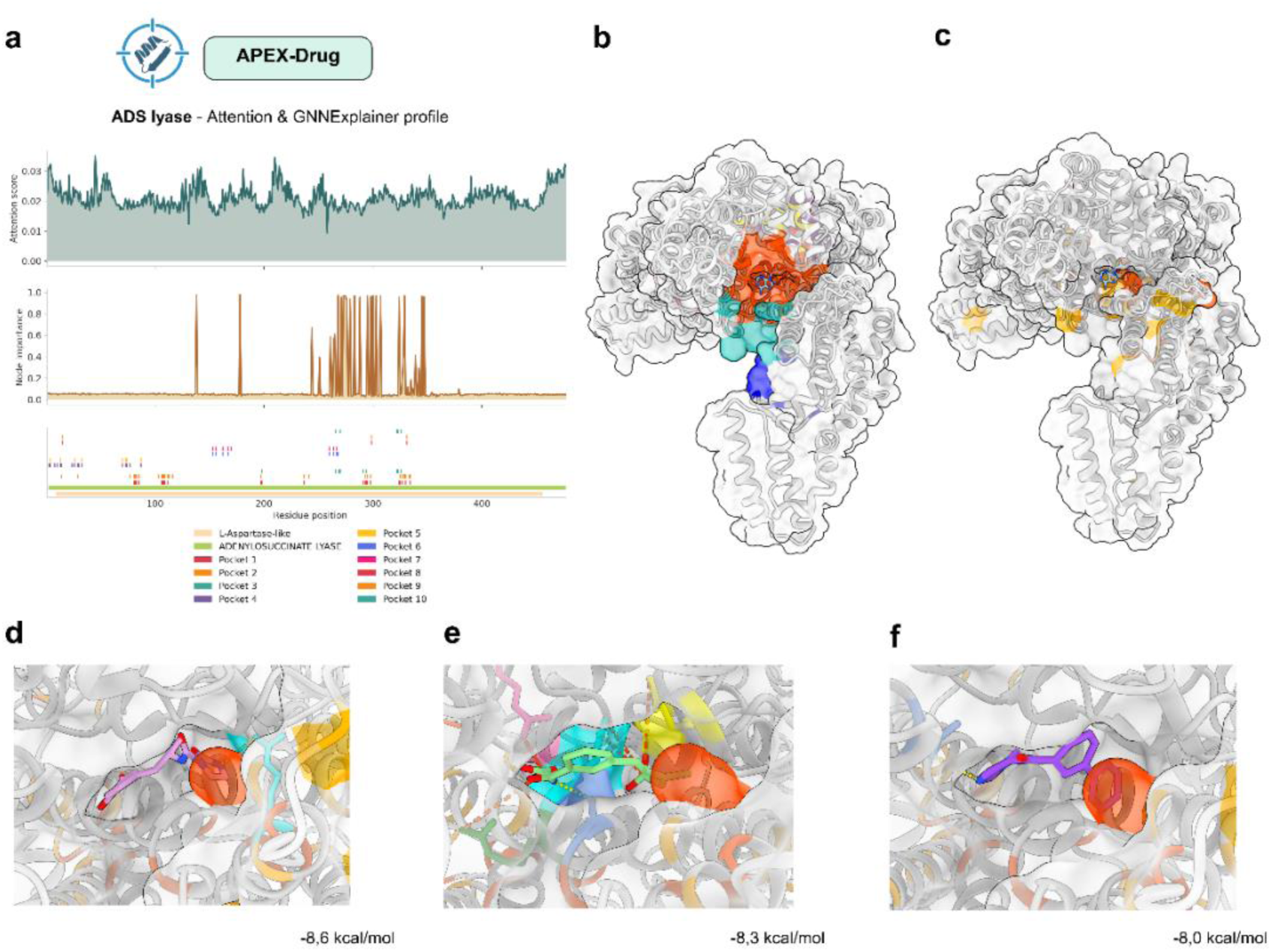
PMDM generates candidate inhibitors for ADSL with predicted strong binding affinity. (a) APEX-Drug interpretability for ADSL showing per-residue attention weights (top, teal) and GNNExplainer importance scores (middle, orange), with Pfam-annotated domains and P2Rank predicted pockets by shown below. (b) AlphaFold3-predicted ADSL structure (dimer, chains B and D) with P2Rank pockets colored and the active site highlighted in orange-red. (c) GNNExplainer-identified important residues mapped onto the structure; the top 10 residues are shown in orange-red, with additional high-ranking residues in orange. (d) Top-ranked PMDM molecule 1 (Vina score −8.6 kcal/mol, pink) forms a hydrogen bond with Arg299-D (3.27 Å, cyan). (e) Second-ranked molecule 2 (Vina score −8.3 kcal/mol, green) forms a hydrogen bond with Ser330-B (2.73 Å, blue) and salt bridges with Arg29-B (3.53 Å, pink), Arg333-B (5.41 Å, forest green), Arg334-B (3.59 Å and 4.90 Å, cyan), and Arg14-D (5.01 Å, yellow). (f) Third-ranked molecule 3 (Vina score −8.0 kcal/mol, purple) forms a hydrogen bond with Thr112-B (3.14 Å). All interacting residues (Ser330-B, Arg29-B, Arg333-B, Arg334-B, Arg14-D, Arg299-D, Thr112-B) lie within the predicted active site. Yellow dashed lines indicate hydrogen bonds; red dashed lines indicate salt bridges.

Using this XAI-defined pocket as a structural constraint, PMDM generated de novo small molecules specifically tailored to the ADSL active site. Of 500 requested molecules, 427 were successfully produced (Supplementary File S5), with a mean docking score of −7.504 kcal/mol. Decoy-based enrichment analysis confirmed that PMDM outputs exhibited substantially higher predicted affinity than physicochemically matched random compounds (ROC-AUC = 0.747; Supplementary File S6). Synthetic feasibility filtering identified 46 synthetically accessible candidates, which were ranked by Vina score and drug-likeness (QED > 0.6), yielding three lead molecules. Molecule 1 (−8.6 kcal/mol) formed a hydrogen bond with Arg299 (3.27 Å) (Figure 5d). Molecule 2 (−8.3 kcal/mol) formed a hydrogen bond with Ser330 (2.73 Å) and salt bridges with Arg29 (3.53 Å) and Arg334 (3.59 Å) (Figure 5e). Molecule 3 (−8.0 kcal/mol) formed a hydrogen bond with Thr112 (3.14 Å) (Figure 5f).

### Pipeline generalization to human bacterial pathogens identifies novel druggable sites in *Acinetobacter baumannii* virulence factors

To test whether APEX generalizes beyond phytopathogenic fungi, we extended the pipeline to human bacterial pathogens. APEX-Tar was retrained on an experimentally validated bacterial virulence-factor dataset, achieving strong performance (AUC = 0.889, AUPR = 0.875, F1 = 0.779, accuracy = 0.819, recall = 0.746, specificity = 0.874) (Figure 6a). Using the same architecture and training strategy, the model learned to distinguish virulence factors from general bacterial proteins based on structural and sequence features encoded in ESM-2 embeddings and residue-contact graphs. We then applied the full prioritization workflow to *Acinetobacter baumannii*, a Gram-negative pathogen classified by the WHO as a critical-priority organism because of its extreme antibiotic resistance. After filtering out homologs to the training dataset, the proteome was evaluated using both bacterial APEX-Tar and APEX-Drug (Supplementary File S4). The top-ranked candidate was the putative fimbrial chaperone YadV, essential for host attachment and biofilm formation (Figure 6b–c).

**Figure 6.**
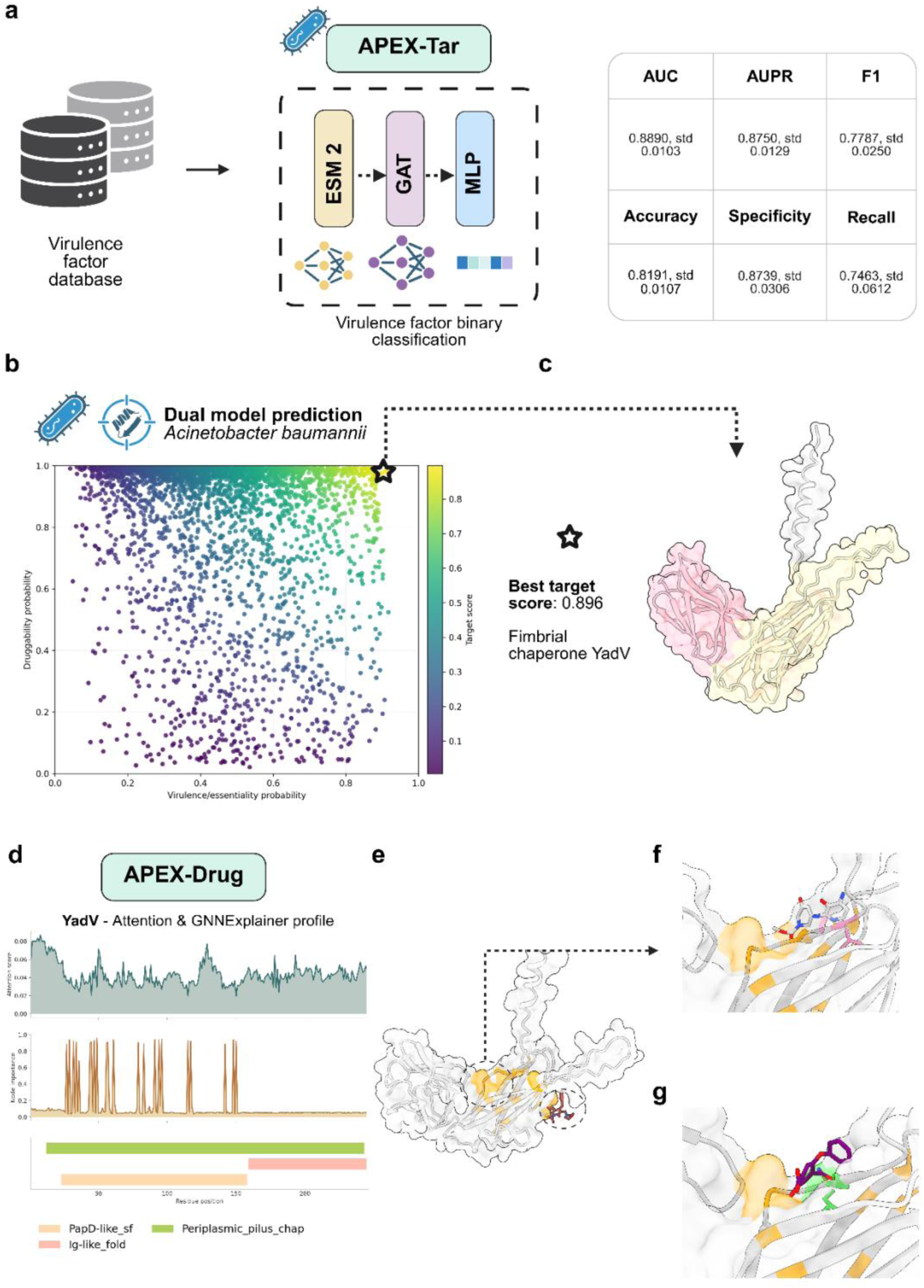
Application of the APEX pipeline to human bacterial pathogens reveals novel druggable sites in *Acinetobacter baumannii* virulence factors. (a) Architecture of the APEX-Tar model trained on the VirulentHunter dataset, achieving an AUC of 0.889, AUPR of 0.875, F1 score of 0.779, accuracy of 0.819, recall of 0.746, and specificity of 0.874. (b) Dual-model predictions for the filtered *A. baumannii* proteome. The scatter plot shows APEX-Tar virulence scores versus APEX-Drug druggability scores, highlighting the fimbrial chaperone YadV as the top combined-score candidate. (c) AlphaFold-predicted structure of YadV showing the characteristic immunoglobulin-like fold of PapD-like chaperones. (d) APEX-Drug interpretability analysis. Attention scores peak at the N-terminal signal peptide, while GNNExplainer identifies key residues within the PapD-like chaperone domain. (e) Validation of the canonical druggable site. Known pilicides (red) dock into the conserved pilin-binding groove, surrounded by GNNExplainer-identified important residues (orange) (f) Novel druggable site predicted by APEX-Drug. PMDM molecule 4 (white; −4.9 kcal/mol) forms a hydrogen bond with Leu29 (light pink; 3.35 Å). (g) Alternative PMDM molecule 5 (purple; −4.4 kcal/mol) targets the same pocket, forming a hydrogen bond with Ile27 (green; 3.18 Å). GNNExplainer nodes (orange) highlight both canonical and newly predicted druggable regions.

APEX-Drug interpretability revealed elevated attention at the N-terminal signal peptide and high GNNExplainer importance within the PapD-like chaperone domain (Figure 6d), the conserved immunoglobulin-like fold characteristic of fimbrial chaperones. Two druggable regions were identified. The first was the canonical pilin-binding groove, previously targeted by pilicides that disrupt chaperone-usher pilus assembly (Figure 6e; Supplementary File S3). Importantly, APEX-Drug also highlighted a second, previously uncharacterized druggable pocket.

PMDM was used to generate small molecules tailored to this novel binding site. Of 500 molecules requested, 315 were successfully generated (Supplementary File S5), with a mean docking score of −4.82 kcal/mol. Decoy-based enrichment analysis confirmed that generated molecules exhibited binding affinity superior to physicochemically matched random compounds (ROC-AUC=0.607; Supplementary File S6). Synthetic feasibility filtering identified 9 synthetically accessible candidates (Supplementary File S5), which were ranked by Vina score and drug-likeness (QED > 0.6), yielding two lead molecules (Figure 6f–g): Molecule 4 showed a predicted binding affinity of −4.9 kcal/mol and formed a hydrogen bond with Leu29 (3.35 Å). Molecule 5 exhibited a predicted affinity of −4.4 kcal/mol and formed a hydrogen bond with Ile27 (3.18 Å).

## Discussion

Identifying new antifungal and antibacterial targets remains difficult across plant and human health, constrained by limited knowledge of pathogen essential genes and increasingly complex resistance mechanisms. Recent AI advances, from protein language models approaching atomic-level structure prediction to generative protein design ^14,15,32^ and machine-learning druggability frameworks ^4,33^, have begun to close these gaps. Yet most methods still treat target discovery and molecular design as separate problems and offer limited mechanistic interpretability.

This study unifies these capabilities by coupling interpretable target discovery with structure-based molecular generation. The pipeline integrates dual GAT models for pathogenicity prediction (APEX-Tar) and druggability assessment (APEX-Drug), target prioritization, XAI-guided site identification, and diffusion-based inhibitor design. Training pathogen-specific APEX-Tar models for both plant-pathogenic fungi and human bacterial pathogens, alongside a universal APEX-Drug model, enables cross-kingdom generalization. This performance suggests that APEX captures fundamental principles of protein function and druggability that transcend taxonomy.

Across benchmarks, all graph-based architectures (GAT, GCN, GraphSAGE) outperformed sequence-only baselines ^25,34,35^, with the largest gains in pathogenicity prediction (APEX-Tar AUC = 0.828 vs. 0.781 ESM-only; Δ=+0.047, p=0.013). Improvements in druggability classification were smaller (APEX-Drug AUROC = 0.961 vs. 0.957), consistent with druggability being driven by conserved structural features already encoded in ESM-2 embeddings. In contrast, pathogenicity involves diverse mechanisms lacking consistent sequence signatures ^36^, making structural context from contact graphs especially valuable.

A key design choice in APEX is retaining evolutionarily related sequences during training. Filtering out proteins above 25–30% identity would remove precisely the conservation signals that define essential and virulence-associated functions. Like protein language model pretraining^14,15,32^, APEX leverages conserved sequence–structure patterns as functional cues rather than treating similarity as a confounder. Sensitivity analysis at a 50% identity threshold confirmed this rationale: APEX-Tar performance changed minimally (ΔAUC = −0.006), indicating reliance on learned evolutionary patterns rather than memorization.

The performance gap between APEX-Tar and APEX-Drug likely reflects differences in annotation quality rather than model architecture. Human druggability labels benefit from decades of pharmaceutical research ^37^, whereas fungal pathogenicity annotations remain heterogeneous across species ^11^. Approaches such as DrugTar highlight the value of combining language model embeddings with functional annotations^33^; our graph-based strategy adds structure-aware learning and residue-level interpretability via GNNExplainer, enabling precise localization of druggable sites for downstream design.

Cross-species transfer learning in APEX-Drug addresses the scarcity of validated targets in non-model pathogens. Training on human proteins and applying the model to fungal and bacterial proteomes exploits the evolutionary conservation of druggability determinants such as pocket geometry, electrostatics, and surface properties ^38^. On a held-out benchmark of pathogen proteins, APEX-Drug correctly classified 96.1%, supporting the idea that small-molecule binding potential is a universal biophysical property ^39^.

Interpretability is a central strength of APEX, distinguishing it from black-box approaches. While post-hoc methods such as SHAP and LIME can be informative, they are computationally heavy and often lack mechanistic clarity ^21^. In contrast, GAT attention weights provide intrinsic interpretability by quantifying information flow during message passing^24^, and GNNExplainer isolates the minimal subgraphs driving predictions ^27^. The alignment of these explanations with known functional sites confirms that the models learn genuine structure–function relationships.

APEX-Tar correctly highlighted the signal peptide and catalytic residues in endopolygalacturonase 1 ^40^, as well as the ATP-binding pocket and MAPK domain in Hog1 ^41^. APEX-Drug recovered benzimidazole and colchicine binding sites in β-tubulin^42^ and the Qo/Qi quinone sites in cytochrome b ^43^.

Manual inspection of top-ranked *B. cinerea* proteins underscored the biological relevance of APEX predictions. Prioritized targets included adenylosuccinate lyase (ADSL), whose *Cryptococcus neoformans* ortholog is essential for virulence ^44^; a SnoaL-like domain protein consistent with roles in toxin biosynthesis ^45^; a deoxyribose-phosphate aldolase involved in nucleotide metabolism ^46^; and enzymes of the β-ketoadipate pathway, a metabolic pathway linked to necrotrophic lifestyles ^47^. Functional enrichment of the 640 high-confidence targets revealed overrepresentation of catalytic activity, carbohydrate metabolism, and secretory pathways hallmarks of necrotrophic fungi that rely on enzymatic host tissue degradation ^44,48^.

Diffusion models complete the target-to-molecule workflow. These models represent the state of the art in structure-based design ^28,29^, generating chemically valid molecules with strong binding potential. Conditioning PMDM on the ADSL active-site geometry produced inhibitors with predicted affinities of −8.6 to −8.0 kcal/mol and favorable drug-likeness. ADSL is an attractive antifungal target due to its role in de novo purine biosynthesis and the presence of fungal-specific active-site residues. Notably, the third-ranked molecule formed a hydrogen bond with Thr112-B, a position occupied by glycine in the human ortholog, suggesting a structural basis for fungal selectivity ^44^. Decoy-based enrichment analysis confirmed that generated molecules outperform physicochemically matched random compounds. While experimental validation remains necessary, the convergence of pocket prediction, interpretability, and docking supports the plausibility of these candidates.

Extending the pipeline to *A. baumannii* further demonstrated its generalizability. Identifying YadV as the top-ranked target highlights an anti-virulence strategy that disrupts colonization without imposing strong selective pressure for resistance ^48^. Fimbrial chaperone-usher pathways are central to adhesion and biofilm formation^49,50^ in *A. baumannii*, and the model recovered both the canonical pilicide-binding groove^51^ and a previously uncharacterized druggable pocket. PMDM-generated inhibitors targeting this novel site formed distinct interactions, including hydrogen bonds with Leu29 and Ile27. Although −4.9 kcal/mol may appear modest, it is comparable to the −5.063 kcal/mol predicted for the canonical pilicide site, suggesting that the newly identified pocket represents a potentially tractable binding site. This result illustrates APEX-Drug’s ability to surface pockets that conventional structure-based workflows might overlook.

Several opportunities exist for refinement. While ESM-2 contact maps enable proteome-scale analysis, incorporating experimental structures or AlphaFold3 multimer predictions could improve modeling of protein–protein and protein–ligand interfaces ^32^. Binary classification simplifies inherently continuous biological properties; although APEX-Drug outputs a continuous druggability probability, experimental validation remains essential. Docking scores provide useful ranking but do not capture entropic, solvation, or conformational effects, and predicted binders may not function as inhibitors if they fail to displace natural substrates or modulate activity. Although PMDM generated chemically valid scaffolds, retrosynthetic analysis identified complete synthetic pathways for only a subset, highlighting synthetic accessibility as a bottleneck. Ligand-based diffusion models may help optimize these scaffolds ^30,52^.

In conclusion, we present an end-to-end computational framework for antimicrobial target discovery and tailored molecular design. The identification of targets in *B. cinerea* and *A. baumannii*, together with de novo inhibitor generation, demonstrates broad applicability across pathogen classes and therapeutic strategies. Built-in interpretability provides mechanistic insight and experimental prioritization, setting APEX apart from black-box approaches. Its modular design makes the framework readily transferable to diverse antimicrobial challenges, offering a scalable blueprint for AI-driven discovery in both agriculture and clinical medicine.

## Methods

### Code availability

The complete APEX pipeline, including training and inference code, is publicly available at https://github.com/Brunxi/APEX. Pre-trained models are available for download at https://zenodo.org/records/18741462.

### Data and preprocessing

Three binary classification datasets were curated for model training. APEX-Tar for phytopathogenic fungi was trained on fungal pathogenicity factors from PHI-base (Pathogen-Host Interactions Database)^53^, selecting proteins annotated with loss of pathogenicity or reduced virulence phenotypes as the positive class (n=1,108) and proteins with unaffected pathogenicity phenotype as the negative class (n=989), yielding 2,097 fungal proteins. APEX-Drug was trained on human protein druggability annotations from the ProTar-II dataset ^33^, comprising 1,221 druggable proteins and 1,124 non-druggable proteins (2,345 total) with experimentally characterized drug-binding properties. APEX-Tar for bacteria was trained on bacterial virulence factors curated from the VirulentHunter dataset^54^, containing 9,148 experimentally validated virulence proteins and 12,184 non-virulence bacterial proteins (21,332 total) from diverse bacterial species. Complete proteomes for genome-wide target prioritization were downloaded from Ensembl: *Botrytis cinerea* B05.10 (assembly ASM83294v1), *Acinetobacter baumannii* (assembly GCA_000963815, ASM96381v1), and *Solanum lycopersicum* (assembly SL3.0). To prevent data leakage across cross-validation folds, sequence redundancy was removed from the full dataset prior to k-fold partitioning using BLASTp^55^ at a 100% sequence identity threshold.

To evaluate the cross-organism transferability of APEX-Drug, druggable proteins from fungal and bacterial pathogens were retrieved from ChEMBL 36 (July 2025) via its public REST API. Twenty species spanning fungi and bacteria were queried, restricting targets to SINGLE PROTEIN entries with a UniProt accession. A target was considered druggable if at least five structurally distinct compounds with pChEMBL ≥ 5.0 (IC50/Ki ≤ 10 µM) had been reported against it. After deduplication, 228 non-human druggable proteins were retained (mean pChEMBL 6.40 ± 0.55), confirming the existence of experimentally validated druggable targets across both kingdoms and providing a held-out benchmark for cross-organism evaluation of APEX-Drug.

Protein sequences were trimmed to a maximum length of 1,000 amino acids to ensure computational tractability. For each sequence, two types of representations were generated using the ESM-2 transformer model (facebook/esm2_t33_650M_UR50D)^15^: per-residue embeddings of 1,280 dimensions obtained from the 33rd layer, and residue-residue contact maps derived from the model’s attention-based contact prediction head. These contact maps encode structural proximity information learned during pre-training on large-scale protein sequence data and serve as the foundation for constructing protein structure graphs.

Protein graphs were constructed by thresholding contact probability maps to identify structurally relevant interactions. For each contact map, a threshold corresponding to the (1-r)th quantile was computed, where r (ratio parameter, default 0.2) controls edge density. Residue pairs with contact probabilities exceeding this threshold were connected by edges, while self-loops were explicitly removed. Residues with no contacts above threshold (isolated nodes) were filtered from the graph to ensure all nodes participate in message passing. The resulting graphs consist of nodes representing individual amino acids with node features given by their 1,280-dimensional ESM-2 embeddings, and edges representing predicted spatial proximity. Edges were duplicated in both directions to create undirected graphs suitable for symmetric message passing operations. This construction preserves a mapping between graph nodes and their original sequence positions, enabling interpretation at the residue level.

### Model architecture and training

A hierarchical Graph Attention Network (GAT) architecture was used ^24^; this architecture processes protein structure graphs through multiple layers of attention-weighted message passing, followed by graph-level pooling and classification. The first GAT layer employs multi-head attention to capture diverse patterns of residue interactions. Given the input node feature matrix *X* ∈ ℝ^*N*×1280^ where N is the number of nodes, each of the 2 attention heads independently computes attention coefficients and aggregates neighborhood information. For a given attention head k and edge (j→i), the attention coefficient is computed as:

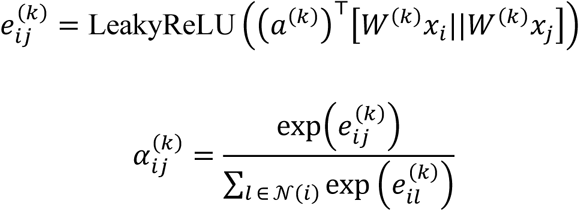

where *W*^(*k*)^ ∈ ℝ^1280×1280^ is a learnable linear transformation, *a*^(*k*)^ ∈ ℝ^2560^ is a learnable attention vector, || denotes concatenation, and *N*(*i*) represents the set of neighbors of node i in the contact graph. The normalized attention coefficients 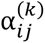 reflect the relative importance of residue j’s features when updating residue i’s representation. The updated feature for node i from head k is:

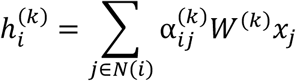

The outputs from all 2 attention heads are concatenated to form 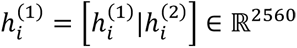, followed by exponential linear unit (ELU) activation: 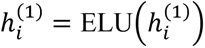. This multi-head architecture allows the model to attend to different structural contexts simultaneously, for example one head focusing on local secondary structure contacts while another captures long-range domain interactions.

The second GAT layer refines the learned representations using a single attention head with the same attention mechanism, transforming the 2,560-dimensional intermediate features back to 1,280 dimensions:

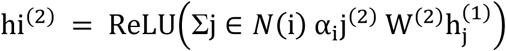

where *W*^(2)^ ∈ ℝ^(2560×1280)^. This layer enables the model to learn higher-order relationships by attending over already-contextualized node representations from Layer 1. Dropout with probability p=0.3 (default) is applied before each GAT layer during training to prevent overfitting, with additional internal dropout on attention coefficients during the attention computation phase to further regularize the learned attention patterns.

Following the two GAT layers, node-level representations 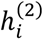 were aggregated into a single graph-level feature vector using global max pooling: 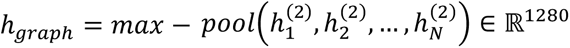. This operation captures the most salient features across all residues, effectively summarizing the entire protein’s learned structural representation. The graph representation is passed through a two-layer fully connected network for binary classification. The first linear layer projects from 1,280 to 16 dimensions with ReLU activation: *z* = *ReLU*(*W*_*fc*1_*h*_*graph*_ + *b*_*fc*1_), *z* ∈ ℝ^16^ The final layer maps to a single output logit: 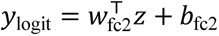, *y*_logit_ ∈ ℝ the predicted probability is obtained via sigmoid transformation: *P*(*y* = 1 ∣ *X*, *E*) = σ(*y*_logit_), where E represents the edge structure. The complete architecture contains approximately 6.6M trainable parameters: 3.3M per GAT layer (dominated by the 1280×1280 weight matrices), plus 20.5K parameters in the classification head. The attention mechanism in each layer adds minimal parameters (attention vectors a) but provides substantial expressive power through learned edge weights.

The dataset was partitioned into 5 folds using stratified k-fold cross-validation (random state = 1029). For each fold, the training set was further split into 80% for training and 20% for validation. Models were trained for 400 epochs using the Adam optimizer with binary cross-entropy with logits loss (BCEWithLogitsLoss). Hyperparameters included learning rate (1×10⁻⁶ for fungi, 7×10⁻⁷ for humans), weight decay (5×10⁻⁴), and batch sizes of 5 for training and 4 for testing. Mixed-precision training with automatic gradient scaling was employed to optimize GPU memory usage. For each fold, minimizing validation loss checkpoints were saved. Final test performance was evaluated using held-out test set for each fold. Two baseline classifiers were evaluated under identical cross-validation splits. The ESM-only baseline used mean-pooled ESM-2 residue embeddings (1,280-dimensional) as input to a logistic regression classifier (L-BFGS, class-weight balanced). The sequence-only baseline used normalized amino acid composition frequencies as features, also classified by logistic regression.

Model performance was assessed using standard binary classification metrics computed at a threshold of 0.5: accuracy, precision (positive predictive value), recall (sensitivity), specificity, F1-score (harmonic mean of precision and recall), area under the ROC curve (AUC), and area under the precision-recall curve (AUPR). Confusion matrices (true positives, false positives, true negatives, false negatives) were computed for detailed error analysis. All metrics were calculated using scikit-learn implementations. For each fold, metrics were reported separately for training, validation, and test sets, with final results aggregated across all 5 folds to provide robust estimates of model performance and uncertainty. Statistical significance of performance differences between GNN architectures and baseline classifiers was assessed using the corrected paired t-test across five cross-validation folds^55^, which accounts for the dependence structure inherent to k-fold evaluation.

To assess the potential impact of residual sequence homology across cross-validation folds in APEX-Tar, sequences were first clustered using CD-HIT v4.8.1 at 50% sequence identity. Cross-fold homologous pairs were then validated by all-vs-all pairwise comparison using MMseqs2 between training and test sets of each fold, retaining the maximum fractional identity per pair. Proteins involved in any cross-fold homologous pair were identified and excluded from test evaluation in a post-hoc sensitivity analysis.

### Model interpretation

To identify residues serving as information integration hubs during classification, attention weights from both GAT layers at inference time were extracted. For each GAT layer, the forward pass yields attention coefficients *α*_*ij*_ for all edges (j→i) in the contact graph. For the multi-head Layer 1, coefficients were averaged across the 2 attention heads. Then, a per-residue attention score was computed by aggregating incoming attention weights:

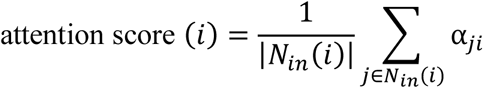

where *N*_*in*_(*i*) represents the set of nodes with edges directed toward node *i*. This metric quantifies how strongly the model weighs information from a residue’s structural neighborhood during message passing. High attention scores indicate residues that act as information integration points, aggregating signals from multiple structural neighbors. Attention scores were computed separately for Layer 1 and Layer 2, then summed to produce a total attention profile across the protein sequence. Residues filtered during graph construction (those with no contacts above threshold) received attention scores of zero. The resulting attention profiles were normalized to the original sequence length to enable comparison across proteins and visualization on linear sequence representations. To identify structurally critical subgraphs necessary for model predictions, GNNExplainer^27^, a perturbation-based explanation method for graph neural networks was applied. GNNExplainer learns a soft mask *M* ∈ [0,1]^*N*^ over nodes by optimizing:

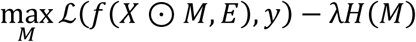

where f represents the trained GAT model, ⊙ denotes element-wise multiplication, E is the edge structure, y is the target class, L is the cross-entropy loss, and H(M) is an entropy regularization term encouraging discrete (binary) masks:*H*(*M*) = − ∑_*i*_[*M*_*i*_ log *M*_*i*_ + (1 − *M*_*i*_) log(1 − *M*_*i*_)]. The hyperparameter λ controls the trade-off between prediction fidelity and mask sparsity. The optimization was performed using Adam (learning rate 0.01, β₁=0.9, β₂=0.999) for 200-400 epochs depending on prediction confidence (400 epochs for predictions with probability >0.95, 200 otherwise).

To validate explanations, insertion-deletion analysis was performed by progressively adding or removing the top-k most important nodes (ranked by learned mask values *M*_*i*_) and measuring the impact on model logits across 11 equally spaced values of k. Four perturbation scenarios were computed: hard removal (extracting the subgraph with top-k nodes and their edges deleted), hard retention (extracting the subgraph containing only top-k nodes and edges among them), soft removal (setting node features to zero for top-k nodes while preserving graph structure), and soft retention (setting node features to zero for all nodes except top-k). Area under the insertion-deletion curves was calculated using the trapezoidal rule to quantify explanation fidelity. Explanations were considered consistent if removing important nodes decreased the predicted class logit or retaining them maintained high prediction scores. By default, the top 10% of nodes (ranked by GNNExplainer mask values) was analyzed as the critical set for detailed interpretation.

### Target prioritization and structural analysis

To identify novel targets, APEX-Tar and APEX-Drug were applied to complete proteomes. Proteins with significant sequence similarity to either training dataset were excluded using BLASTp (BLAST+ version 2.12.0+)^56^ with parameters: e-value threshold 1×10⁻⁵, minimum sequence identity 50%, and minimum query coverage 50%. This filtering ensures the pipeline discovers genuinely novel biological insights rather than recognizing training examples by sequence homology. For fungal targets, proteins with homologs in the *S. lycopersicum* proteome were additionally removed using identical BLASTp criteria to minimize potential off-target effects on host proteins and reduce phytotoxicity risk. Target candidates were ranked by a combined target score defined as the product of APEX-Tar and APEX-Drug prediction probabilities: Target Score = P(pathogenicity/virulence) × P(druggability). High-confidence targets were selected by applying stringent dual thresholds (P > 0.6 for both models). This combined score serves as a heuristic prioritization metric rather than a calibrated joint probability. To assess the independence assumption implicit in the product formulation, we computed the Spearman correlation between APEX-Drug and APEX-Tar output scores across the full proteomes of both organisms, yielding ρ = −0.199 (n = 8,042) for *B. cinerea* and ρ = −0.195 (n = 3,102) for *A. baumannii*. The weak negative correlation confirms that the two models capture largely orthogonal biological features (druggability and pathogenic essentiality are not redundant signals) supporting the use of their product as a composite ranking criterion.

Three-dimensional protein structures were retrieved from the AlphaFold Protein Structure Database^13^, which provides pre-computed high-accuracy structure predictions for reference proteomes. For proteins not available in AlphaFold DB, structures were predicted using Alphafold2^14^. Protein domain annotations and functional sites were obtained from InterPro^57^ by submitting amino acid sequences to the web interface. Signal peptides were predicted using SignalP 6.0^58^. Structural pockets were predicted using the P2Rank web server ^59^. AlphaFold-predicted structures were submitted in PDB format with default server parameters.

Gene Ontology (GO) term enrichment analysis was performed using DAVID (Database for Annotation, Visualization and Integrated Discovery)^60^ web server. Prioritized target protein identifiers were submitted as protein lists with the corresponding complete proteome as background reference. Enriched GO terms for Biological Process, Molecular Function, and Cellular Component categories were identified using Fisher’s exact test with Benjamini-Hochberg false discovery rate (FDR) correction. Terms with adjusted p-value < 0.05 were considered significantly enriched and reported.

### Structure-based molecular design

Novel small-molecule inhibitors were generated using PMDM (Pocket-based Molecular Diffusion Model) ^31^with the following configuration: 3D coordinates of residues within the predicted binding pocket as input, maximum 20 atoms per molecule, 500 molecules generated per target, batch size 1, and GPU acceleration enabled. The 250 molecules with the most favorable docking scores were retained for downstream analysis. Characterized ligands (pilicides for YadV^61^, benzimidazoles for β-tubulin^42^, strobilurins for cytochrome b^62^ and AMP for ADSL ^44^) were docked using DiffDock^63^. Binding affinities for PMDM-generated molecules were predicted using QVina2.1 (exhaustiveness=16, box size=20 Å per axis), with the docking box centered on the predicted pocket centroid ^64^. Binding energies were reported as Vina scores (kcal/mol).

To assess whether generated molecules exhibit binding affinity superior to random drug-like compounds, physicochemically matched decoys were generated using the LUDe v1.0 ChEMBL30 database^65^, selecting candidates within ±20 Da molecular weight, ±0.5 logP, ±1 hydrogen bond acceptors/donors, ±1 rotatable bonds, and identical formal charge. Dissimilarity was enforced by retaining only decoys with Tanimoto similarity ≤0.2 (Morgan fingerprints, radius=2, 2048 bits). Up to 250 decoys per target were docked under identical conditions. Enrichment was quantified by ROC-AUC and enrichment factor at 1% (EF1%).

Hydrogen bonds, salt bridges, and hydrophobic contacts were characterized using BINANA^66^ with geometric criteria: hydrogen bonds (donor-acceptor distance ≤ 3.5 Å, donor-hydrogen-acceptor angle ≥ 120°), salt bridges (oppositely charged atoms within 5.5 Å), and hydrophobic contacts (carbon-carbon distance ≤ 4.0 Å).

Molecular properties were calculated using RDKit version 2023.09.1: QED (Quantitative Estimate of Drug-likeness, 0-1 scale)^67^, SA Score (Synthetic Accessibility, 1-10 scale)^68^, logP^69^, and Lipinski’s Rule of Five compliance (molecular weight ≤ 500 Da, logP ≤ 5, hydrogen bond donors ≤ 5, hydrogen bond acceptors ≤ 10)^70^. Molecules were exported in SMILES format. Synthetic feasibility was evaluated using AiZynthFinder 4.0^71^ model (time limit=900 s/molecule, 150 iterations, maximum 10 transforms per step), using USPTO and RingBreaker retrosynthesis models against a ZINC commercial stock database. Molecules for which AiZynthFinder identified a complete retrosynthetic route to commercially available building blocks were classified as synthetically feasible.

### Implementation details

All models were implemented in PyTorch 2.7.0^72^ with PyTorch Geometric 2.7.0^73^ for graph operations. ESM-2 models were loaded via the fair-esm library (version 2.0.0)^15^. Training was performed on NVIDIA DGX Station equipped with 4× Tesla V100-DGXS-32GB GPUs (32GB VRAM each) with CUDA 13.0 support. Additional dependencies included NumPy 2.0.2, pandas 2.2.3, matplotlib 3.9.4, BioPython 1.86, and tqdm 4.67.3. Experiments were optionally tracked using Weights & Biases 0.23.1. For reproducibility, random seeds were set to 1029 for data splitting and 42 for explanation methods. All computations were executed on the Picasso HPC cluster at SCBI-UMA.

### Language model assistance

Large language models were used to assist with manuscript preparation. Claude 4.6 Sonnet (Anthropic, San Francisco, CA), ChatGPT 5.4 (OpenAI, San Francisco, CA) and Copilot (Microsoft, Redmond, WA) were employed for English language translation and text synthesis to improve clarity and readability. All scientific content, experimental design, data analysis, interpretation of results, and conclusions were generated solely by the authors. The authors take full responsibility for the accuracy and integrity of the work.

## Author contributions

Á.P and L.J-C. designed and planned the experiments. L.J.C. performed the experiments. L.J.C, A.P-G and Á.P. wrote the manuscripts. A.P-G., D.F-O and Á.P. revised the manuscript. Á.P. supervised the study. All authors read and approved of its content.

